# A morphological and molecular analysis of the species diversity of the cichlid genus *Petrochromis* from Lake Tanganyika (Teleostei: Cichlidae)

**DOI:** 10.1101/280263

**Authors:** Carl Mattsson

## Abstract

A taxonomic revision of the cichlid fish genus *Petrochromis* endemic to Lake Tanganyika. All recognized taxa are herein described, one subspecies is given species status and five new species, viz. *P. calliris, P. daidali, P. heffalumpus, P. lisachisme* and *P. paucispinis*, are presented. *P. calliris* is described from 6 specimens from Cape Mpimbwe, and distinguished primarily by having a high number of both gill rakers on the lower limb of the first gill arch and vertebrae. *P. daidali* is described from 18 specimens from Cape Mpimbwe, Kansombo and Nkwazi point, and distinguished primarily by males having a labyrinth-like pattern on the head. *P. heffalumpus* is described from 7 specimens from Cape Mpimbwe, and distinguished primarily by its great size. *P. lisachisme* is described from 12 specimens from Cape Mpimbwe and Lyamembe, and distinguished primarily by having a high number of dorsal spines. *P. paucispinis* is described from 4 specimens from Halembe, and distinguished primarily by having a low number of dorsal spines. A revised key to *Petrochromis* is included. A phylogenetic tree hypothesis of the genus, based on molecular (mitochondrial cytochrome *b* and d-loop) and morphological results show that jaw position and number of vertebrae are important diagnostic characters. Analyses suggest that the “ancestral *Petrochromis*” might have looked something like *P. orthognathus*.

## Introduction

Lake Tanganyika (Fig. 1), the 0oldest of the Rift Valley lakes in east Africa, perhaps as old as 20 million years (Brichard, 1989), is the second largest freshwater lake in the world by volume and also the second deepest after Lake Baikal in Siberia (Ndembwike, 2006). It is one of the most important biodiversity hotspots in the world with an almost completely endemic lake fauna (Marijnissen, et al. 2006). Among the cichlids in the lake, the algal grazing niche is the most diverse and also has the highest number of species (Kolm, 2009).

**Fig. 1.**
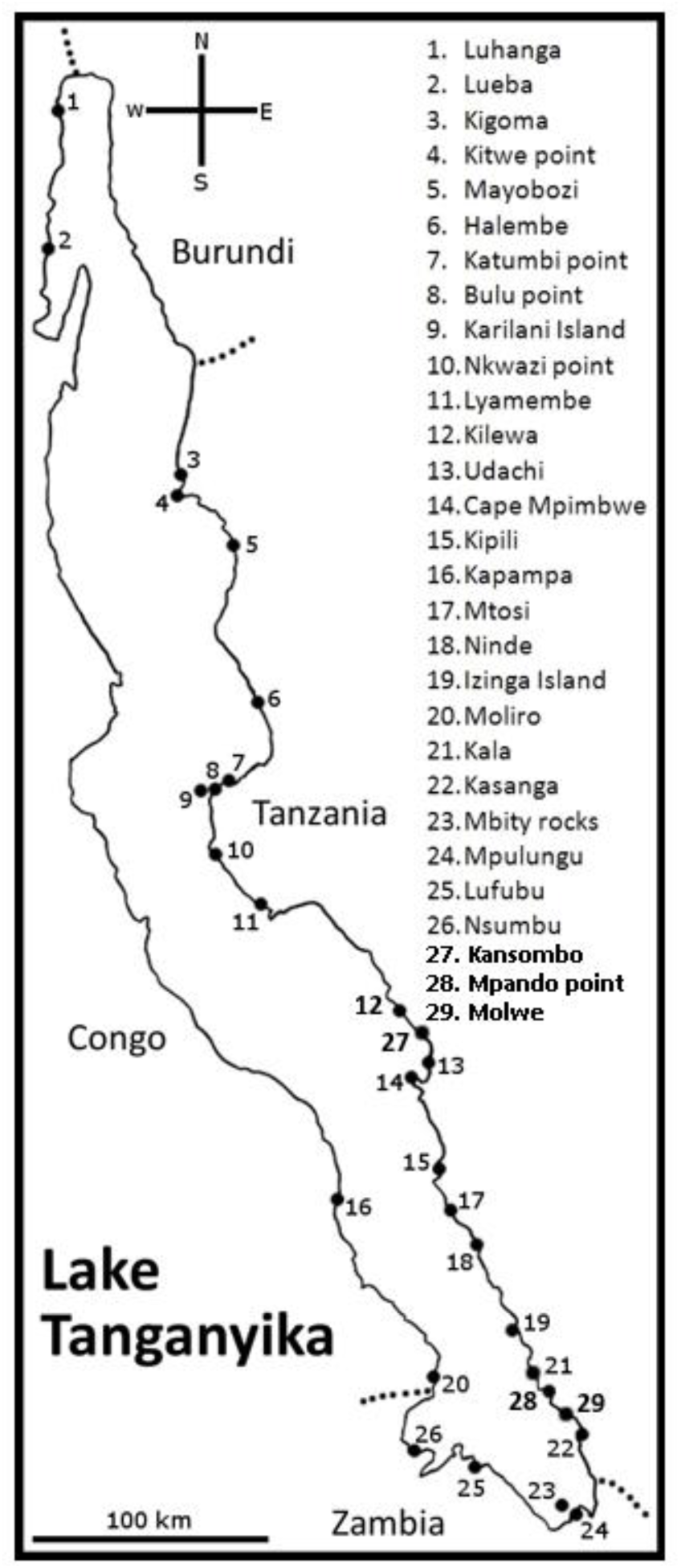
Map of Lake Tanganyika.

*Petrochromis* is a genus of cichlid fishes endemic to Lake Tanganyika in East Africa (Yamaoka, 1983). They are highly specialized algal grazers that feed by scraping their teeth against rocks covered with epilithic growth (Yamaoka, 1983). *Petrochromis* inhabit shallow rocky shores (Yamaoka, 1983) but there is evidence to suggest that there are several species living in deeper water (Konings, 1996). *Petrochromis* are noted for their large size and for their aggression compared to other cichlids in the lake (Brichard, 1989).

*Petrochromis* was established by Boulenger (1898) with *P. polyodon* as the only species. Up till now six species and one subspecies have been described (Brichard, 1989). *Petrochromis polyodon* was based on four specimens from Kinyamkolo (presently Mpulungu) and Mbity Rocks off the southernmost coast of Lake Tanganyika (Boulenger, 1898). Boulenger (1902) described a new species, *P. nyassae*, based on a single specimen from Lake Nyassa (presently Lake Malawi). It was later found to be a specimen of *P. polyodon* with erroneous locality data as discussed by Matthes & Trewavas (1960). The second valid species in the genus was described by Boulenger (1914) as *P. fasciolatus*, based on six specimens from Kapampa on the western shore and Kilewa Bay on the eastern shore of Lake Tanganyika. It was diagnosed by having numerous vertical stripes on its body (Boulenger, 1914). Poll (1948) described the third valid species, *P. trewavasae*, based on a single specimen from Moliro off the southwest coast of Lake Tanganyika, diagnosed by its lunate caudal fin. Matthes (1959) described the fourth valid species, *P. orthognathus*, based on 26 specimens from the northwest part of Lake Tanganyika, from Luhanga in the north to Lueba in the south. This species was diagnosed primarily by its isognathous jaws. The fifth valid species, *P. famula*, was described by Matthes & Trewavas (1960) based on 37 specimens from different localities in Lake Tanganyika. The localities stretched all across the lake from Luhanga in the northwest to Kasanga in the southeast. *Petrochromis famula* was diagnosed primarily by its slightly isognathous jaws and also the lack of prominent cheek scales. Yamaoka (1983) described the sixth valid species, *P. macrognathus*, based on six specimens from Luhanga in the northwest part of Lake Tanganyika. It was diagnosed primarily by its projecting upper jaw. Brichard (1989) described a new subspecies of*P. trewavasae* called *P. trewavasae ephippium*. Type information and locality was not given in the original description. It was diagnosed by not having as long and filamentous fins as *P. t. trewavasae*.

Farías (2001) states that cytochrome *b* is useful when assessing relationships between East African cichlids and Sturmbauer (2003) used both this gene and the gene d-loop in his molecular study on the tribe *Tropheini*, which includes e.g. *Petrochromis* and *Tropheus*. Since Sturmbauer (2003) only partially resolved the phylogeny of *Petrochromis* (Fig. 2) the present paper includes both molecular and morphological analyses for this purpose.A revision of the genus is in order since several new species have been collected, notably some deep water species (Table 1), and the objective of the present paper is to review existing taxa, describe new species available, provide a revised key to the genus (Table 6) and provide a first relationship hypothesis (Fig. 7) for the expanded set of species.

**Fig. 2.**
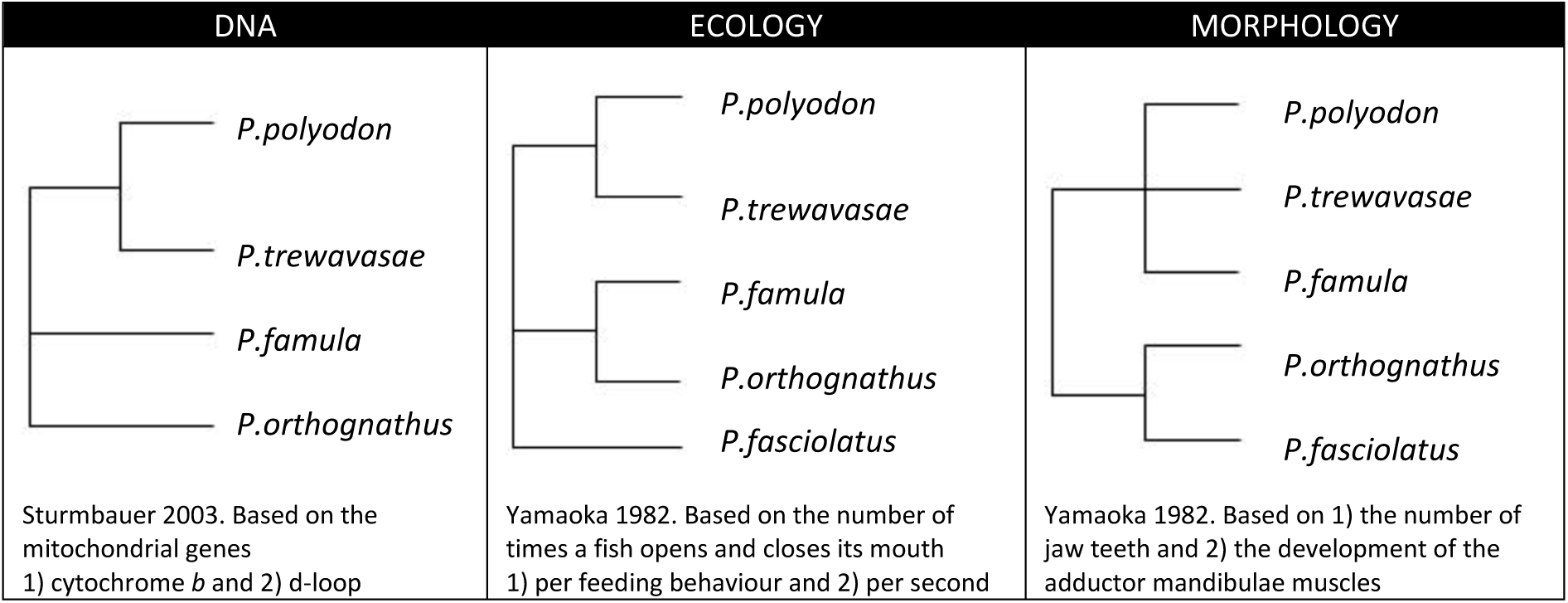
Earlier hypotheses of the phylogeny of species of *Petrochromis*.

**Table 1.**
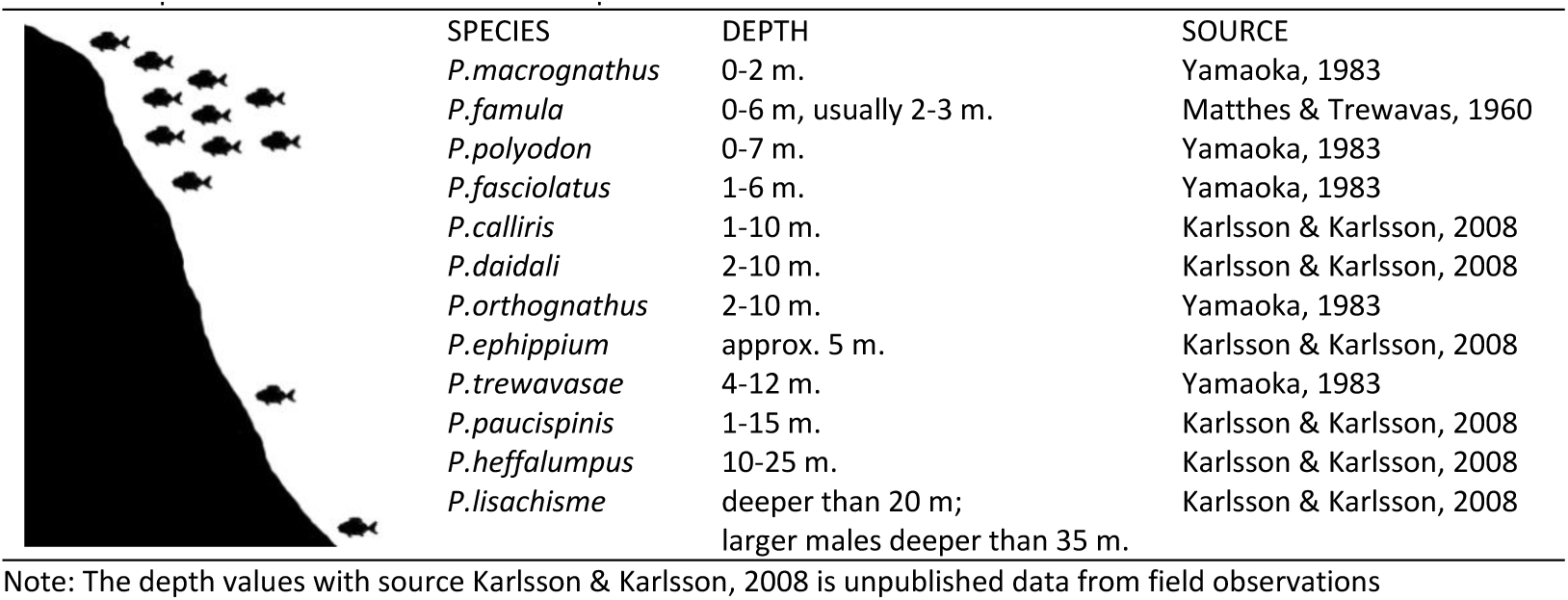
Depth distribution of *Petrochromis* species

|  | SPECIES | DEPTH | SOURCE |
| --- | --- | --- | --- |
|  | *P. macrognathus* | 0-2 m. | Yamaoka, 1983 |
|  | *P. famula* | 0-6 m, usually 2-3 m. | Matthes & Trewavas, 1960 |
|  | *P. polyodon* | 0-7 m. | Yamaoka, 1983 |
|  | *P. fasciolatus* | 1-6 m. | Yamaoka, 1983 |
|  | *P. calliris* | 1-10 m. | Karlsson & Karlsson, 2008 |
|  | *P. daidali* | 2-10 m. | Karlsson & Karlsson, 2008 |
|  | *P. orthognathus* | 2-10 m. | Yamaoka, 1983 |
|  | *P. ephippium* | approx. 5 m. | Karlsson & Karlsson, 2008 |
|  | *P. trewavasae* | 4-12 m. | Yamaoka, 1983 |
|  | *P. paucispinis* | 1-15 m. | Karlsson & Karlsson, 2008 |
|  | *P. heffalumpus* | 10-25 m. | Karlsson & Karlsson, 2008 |
|  | *P. lisachisme* | deeper than 20 m;<br>larger males deeper than 35 m. | Karlsson & Karlsson, 2008 |

Note: The depth values with source Karlsson & Karlsson, 2008 is unpublished data from field observations

## Materials & Methods

### Molecular analysis

A total of 67 mitochondrial DNA sequences (28 cytochrome *b* and 39 d-loop) were used for molecular phylogenetic analyses (Table 2). Complete mitochondrial cytochrome *b* sequences (1137 bp) were obtained from 12 specimens of *Petrochromis*, including the potentially new species *P*. sp. Rainbow Kipili and *P*. sp. Yellow, sampled from live aquarium specimens and processed as fresh tissue; and from two specimens of *Tropheus* preserved in ethanol. Additional sequences of cytochrome *b* and the mitochondrial d-loop (338 bp) were downloaded from the NCBI GenBank (http://www.ncbi.nlm.nih.gov/Genbank/). Those cytochrome *b* sequences are only 402 bp long, and the dataset adjusted accordingly.

**Table 2.**
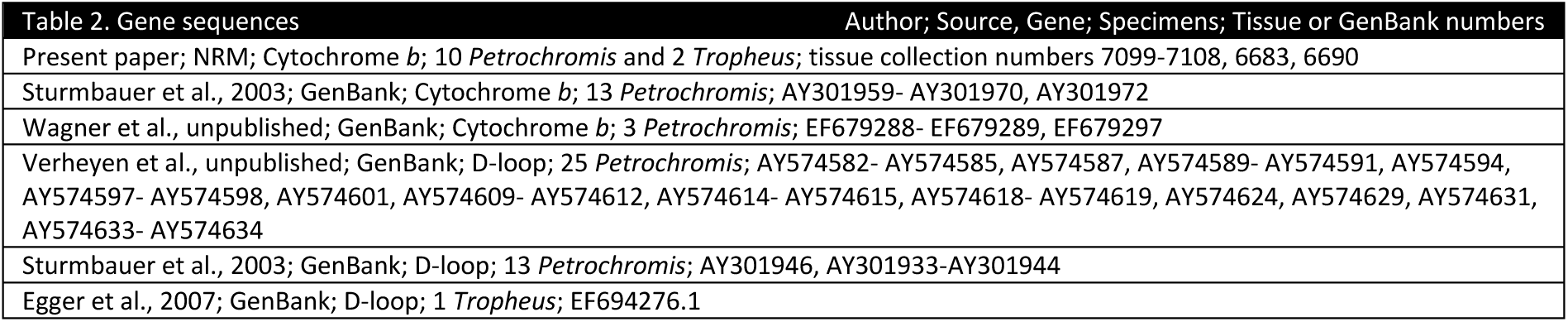
Genesequences

DNA was extracted, using MoleStrips DNA tissue kit (Mole Genetics), with 100 μl lysis buffer and 10 μl proteinase K (conc. 10 mg/ml), and then processed in a GeneMole automated DNA extraction instrument (Mole Genetics). A PCR was then performed using Illustra hot start mix RTG (GE Healthcare) and a final primer concentration of 0.2 μM. The following primers were used: FishCytB-F (ACCACCGTTGTTATTCAACTACAAGAAC); TrucCytB-R (CCGACTTCCGGATTACAAGACCG); CytBi5R (GGTCTTTGTAGGAGAAGTATGGGTGGAA); CytBi7F (CTAACCCGATTCTTTGCCTTCCACTTCCT). Each reaction contained 4 μl of DNA and 0.5 μl of each primer. Sterile distilled water was added to give a final reaction volume of 25 μl. A touchdown PCR program started with initial denaturation at 94°C for 4 min, followed by four cycles of 94 °C for 30 s, 55 °C for 30 s, 72 °C for 1 min, and another four-cycle phase and one 35-cycle phase with identical temperatures and intervals, except that the annealing temperatures were reduced to 53 °C and 51 °C respectively. The program ended with 72 °C for 8 min. Enzymatic PCR product clean-up was then done with Exonuclease I, *E. Coli* (Fermentas). Each PCR reaction mixture contained Exonuclease (20 u) and FastAP thermosensitive alkaline phosphatase (4 u). Cycle sequencing was performed using the ABI BigDye kit (Applied Biosystems) and the products were cleaned using DyeEx kit (Qiagen). The fragments were separated on an ABI 3130xl Genetic Analyzer (Applied Biosystems). Sequences were then assembled and checked with the Staden software (Staden & Bonfield, 1998) and aligned manually with the BioEdit software (Hall, 1999). Regions of partially incomplete data at the beginning and end of the sequences were identified and excluded from subsequent analysis. Using the computer shell program PaupUp 1.0.3.1 (Calendini & Martin, 2007), incorporating the software PAUP* 4.0 (Swofford, 2000), Treeview 1.6.6 (Page, 2001) and Modeltest3.7 (Posada, 2008), a Maximum Likelihood analysis was performed on the data from d-loop and cytochrome *b* respectively. Tree support was calculated using the bootstrap procedure, with 1.000 replicates, in PAUP*4.0 (Swofford, 2000). Then a Bayesian analysis of the two genes was also made using MrBayes 3.1.2 (Ronquist, 1991). *Tropheus* was used as outgroup for tree rooting.

### Morphological analysis

All in all 14 measurements and 7 meristic counts (fin rays, scales & vertebrae) were taken (Tables 4-5) from a total of 145 specimens. Measurements from 6 juveniles and 17 aquarium specimens were excluded from analysis. Methods for taking counts and measurements follow Barel et al. (1977) except for the following; pectoral fin length (from the tip of the longest fin ray to the base of the uppermost fin ray), length of lower jaw (from the anterior tip to the posterior end of the lower jaw), length of upper jaw (from the anterior tip to the posterior end of the upper jaw), labial depth (the deepest part of the upper lip), length of caudal peduncle (from the posterior measuring point of the standard length to the base of the last anal-fin ray) and length of last dorsal spine (from the tip to the base of the last dorsal spine). Vertebral counts were obtained from radiographs.

Seventeen characters were selected for morphological phylogenetic analysis, using Nexus Data Editor 0.5.0 (Page, 2001) to create the data matrix (Fig. 3, Table 3). The resulting Nexus file was then analyzed using PAUP*4.0 (Swofford, 2000) employing the heuristic tree search algorithm. *Tropheus* was used as outgroup for tree rooting. Tree support was calculated using the bootstrap procedure, with 10.000 replicates, in PAUP*4.0 (Swofford, 2000).

**Fig. 3.**
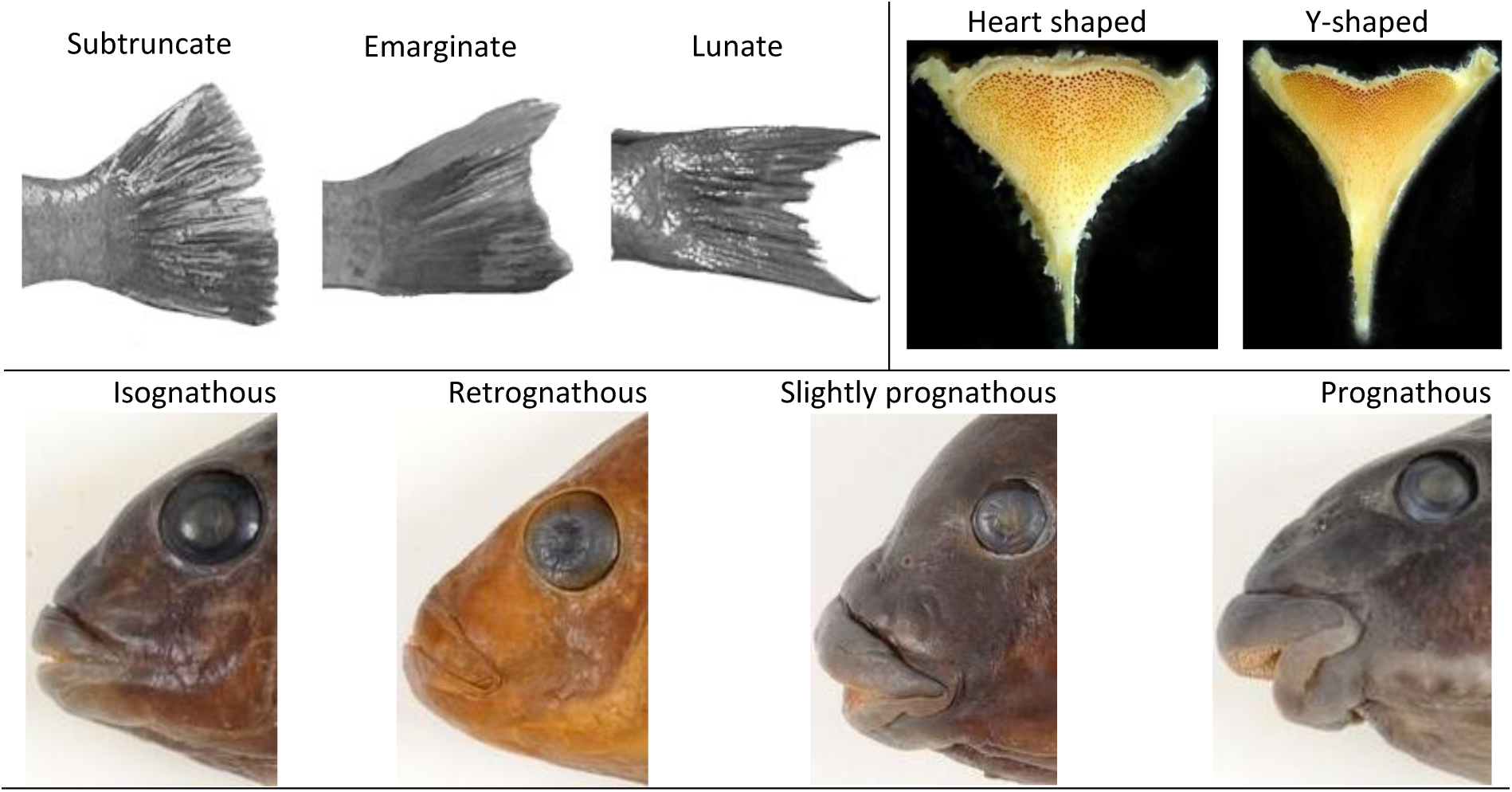
Shape of caudal fin, lower pharyngeal jaw bone and upper jaw position in species of *Petrochromis*.

**Fig. 4.**
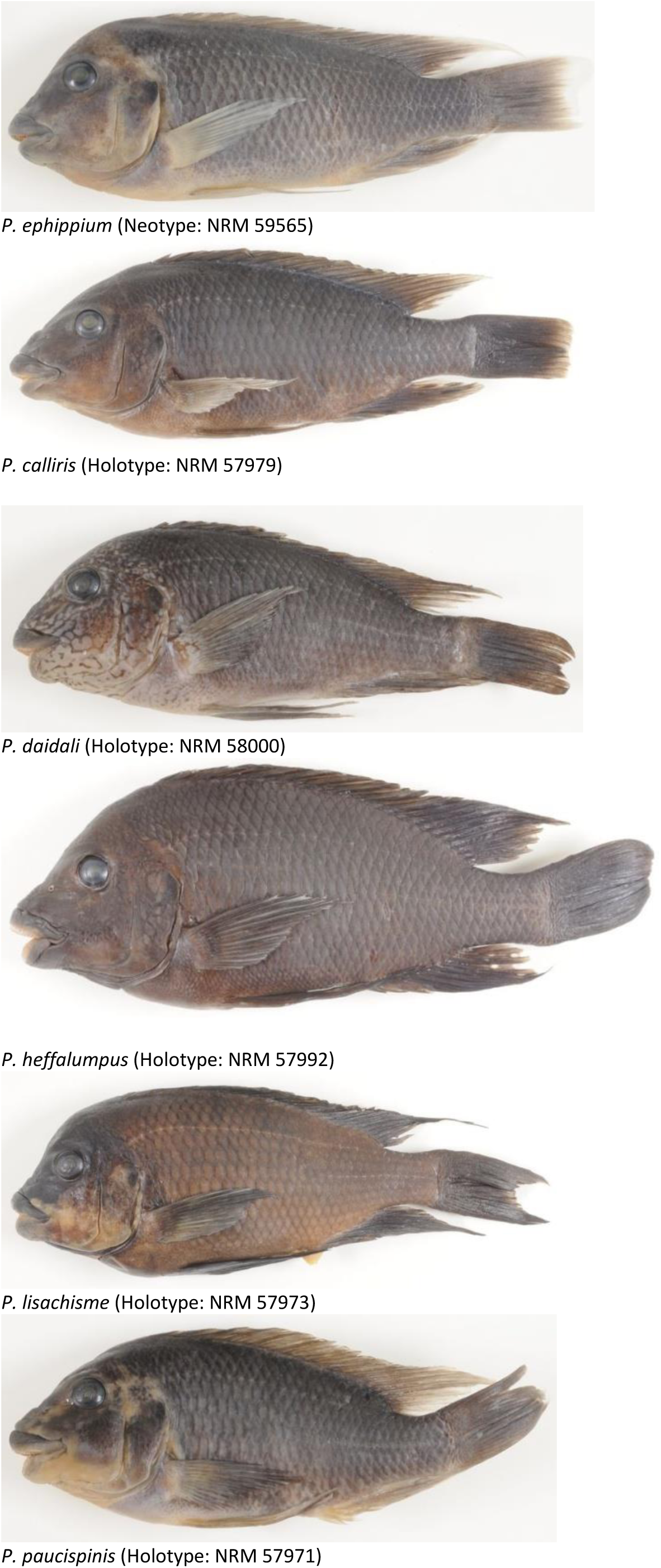
Holotypes and live color photographs of *Petrochromis*; and neotype of *P. ephippium*.

**Table 3.**
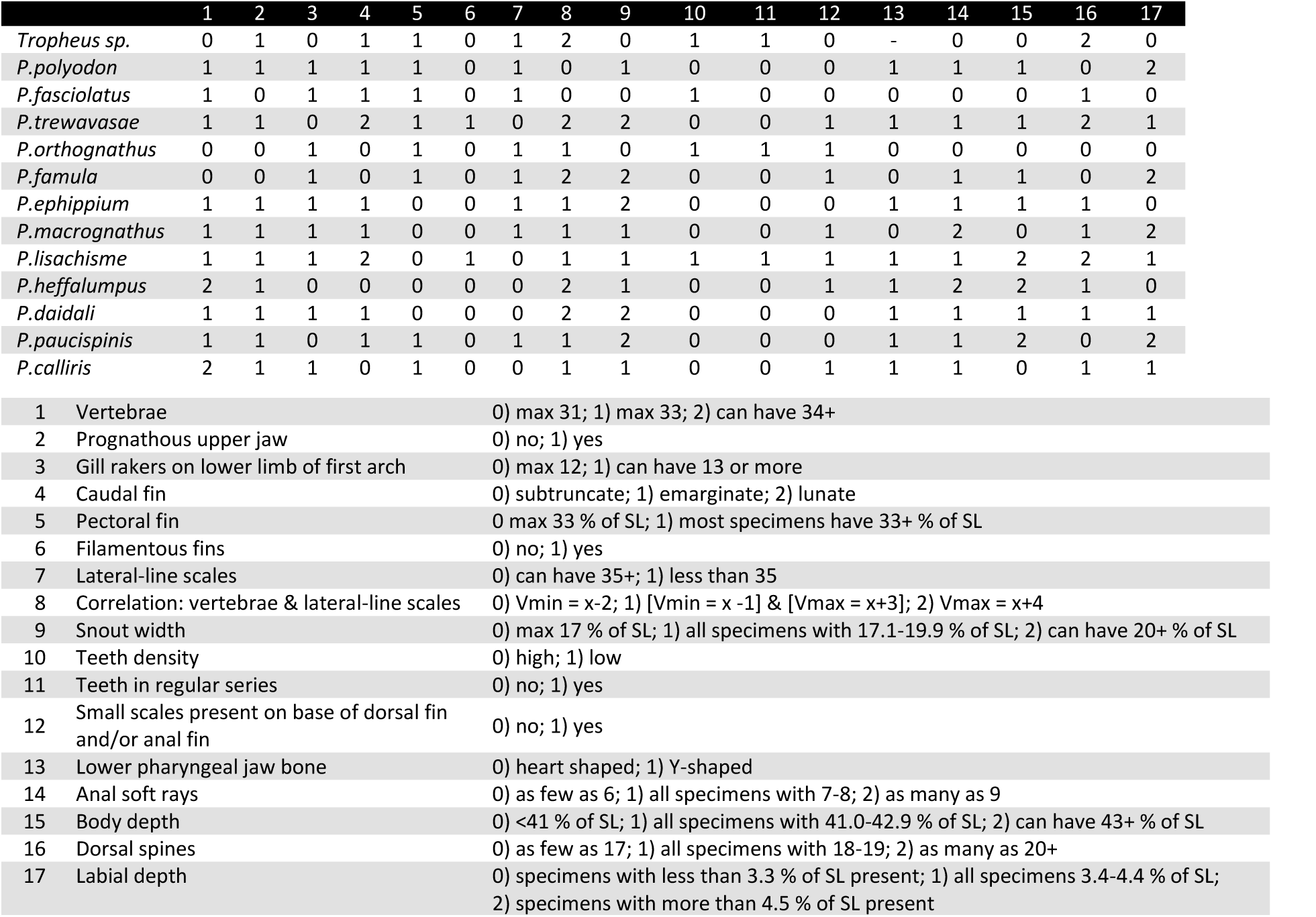
Data matrix based on morphological features of *Petrochromis* and *Tropheus*

Specimens are preserved in the fish collection in the Swedish Museum of Natural History, Stockholm (NRM). Specimens, locality information, ornamental fish trade names, data on habitat and feeding behaviour of the five new species were provided by Magnus and Mikael Karlsson (in litt.) of the African Diving LTD.

## Results

### Genus ***Petrochromis*** Boulenger, 1898

**Diagnosis:** Distinguished from other cichlids by having dental pads with a large number of tricuspid teeth on slender shafts (Yamaoka, 1983).

**Description:** Dorsal fin XVII-XXI, 8-11; anal fin III, 6-9; pectoral fin 13-15; number of lateral-line scales 30-38; vertebrae 30-35; gill-rakers on lower limb of first gill arch 10-15. Compact body and wide snout. Mouth large. Lips thick. Many tricuspid teeth with slender shafts in the jaws. Pelvic fins pointed with the first soft ray the longest. Egg dummies present on soft dorsal and anal fins. Caudal fin emarginate, subtruncate or lunate. Only a right gonad present.

### ***Petrochromis polyodon*** Boulenger, 1898

**Diagnosis**: D. XVII-XIX, 8-10; A. III, 7-8; P. 13-14; gill-rakers on lower limb of first gill arch 12-14; lateral-line scales 30-34; vertebrae 31-33 (usually 32). **Material examined**: NRM 12884, 2 males, 113.3-137.0 mm SL, Congo, Luhanga. NRM 24157, 4 females and 1 male, 42.0-63.9 mm SL, Zambia, fishing camp midway between Lufubu and Mpulungu. NRM 51317, female, 170.3 mm SL, Tanzania, Molwe. NRM 60210, 6 males, 125.6-162.8 mm SL, Tanzania, Mpando point.

### ***Petrochromis fasciolatus*** Boulenger, 1914

**Diagnosis**: D. XVIII-XIX, 8-10; A. III, 6-8; P. 13-14; gill-rakers on lower limb of first gill arch 11-13; lateral-line scales 30-34; vertebrae 31-33 (usually 32). **Material examined**: NRM 12757, male, 88.5 mm SL; NRM 12825, 2 males, 28.8-36.8 mm SL; Zambia, Nsumbu. NRM 12888, 2 males, 86.7-99.6 mm SL, Congo, Luhanga. NRM 51525, male, 103.3 mm SL, Tanzania, Udachi. NRM 51286, 7 males and 3 females, 103.1-124.6 mm SL; NRM 60215, female, 106.2 mm SL; Tanzania, Kansombo.

### ***Petrochromis trewavasae*** Poll, 1948

**Diagnosis**: D. XIX-XX, 8-9; A. III, 7-8; P. 13-14; gill-rakers on lower limb of first gill arch 10-11; lateral-line scales 32-36; vertebrae 31-32. **Material examined**: NRM 13300, 8 females and 5 males, 73.2-105.0 mm SL, aquarium specimens. NRM 16893, 3 females, 105.1-128.2 mm SL, Tanzania, Kitwe point. NRM 17434, male, 30.5 mm SL, Tanzania, Kigoma.

### ***Petrochromis orthognathus*** Matthes, 1959

**Diagnosis**: D. XVIII-XIX, 8-9; A. III, 6-8; P. 13-14; gill-rakers on lower limb of first gill arch 11-15; lateral-line scales 30-33; vertebrae 31. **Material examined**: NRM 17447, male, 118.0 mm SL, Tanzania, Kigoma. NRM 33695, 3 males and 1 female, 71.7-101.5 mm SL, aquarium specimens.NRM 51321, 10males, 99.2-128.2 mm SL; NRM 57987, 8 males and 2 females, 93.0-109.4 mm SL; NRM 60213, male,122.2 mm SL; Tanzania, Cape Mpimbwe.

### ***Petrochromis famula*** Matthes & Trewavas, 1960

**Diagnosis**: D. XVII-XVIII, 8-10; A. III, 8; P. 13-14; gill-rakers on lower limb of first gill arch 11-14; lateral-line scales 32-34; vertebrae 30-31 (usually 30). **Material examined**: NRM 51322, 6 females and 2 males, 97.3-136.2 mm SL; NRM 60214, male, 123.9 mm SL; Tanzania, Cape Mpimbwe.

### ***Petrochromis macrognathus*** Yamaoka, 1983

**Diagnosis**: D. XVIII, 10; A. III, 7-9; P. 15; gill-rakers on lower limb of first gill arch 12-14; lateral-line scales 34; vertebrae 32-33 (usually 33). **Material examined**: NRM 51511, male, 89.8 mm SL, Tanzania, Molwe. NRM 59631, 4 males and 1 female, 137.3-150.0 mm SL, Tanzania, Katumbi point.

### ***Petrochromis ephippium*** Brichard, 1989

**Diagnosis**: D. XIX, 9; A. III, 7; P. 13-14; gill-rakers on lower limb of first gill arch 12-14; lateral-line scales 32-33; vertebrae 31-33 (usually 32). **Material examined**: NRM 51320, male, 152.1 mm SL; NRM 59565, 5 males and 2 females, 127.5-156.9 mm SL; Tanzania, Udachi, 2 Feb 2008, M. Karlsson.

**Remarks**: Brichard (1989) described “*P. t. ephippium*” as a subspecies to *P. trewavasae* noting three major differences from the nominate subspecies; 1) not having as long and filamentous fins; 2) color brown to yellow-orange, instead of dark brown with white spots; 3) the geographical distribution of the two, with *P.trewavasae* restricted to a small coastal area on the west coast, approximately from Moliro to Kapampa, and that “*P. t. ephippium*”, being more well spread, had only been found some way north of that and also in the southern part of the lake. Since I have examined east coast specimens of both of these species the geographical notion can be disregarded.

Brichard (1989) could find no significant morphometric differences between *P. ephippium* and *P. trewavasae* but *P. ephippium* has an emarginate caudal fin and 12-14 gill rakers on the lower limb of the first arch whereas *P. trewavasae* has a lunate caudal fin and 10-11 gill rakers.

*Petrochromis ephippium* is referred to in the ornamental fish trade as *P.* sp. Moshi. This I have deducted from studying live aquarium specimens and it being described as having a body color of brown to yellow-orange (Konings, 1988; Herrmann, 1985; Nemeth, 2009 unpubl.) and sometimes a saddle-like rectangular light-coloured patch under the dorsal fin (Brichard, 1989; Herrmann, 1985).

## Five new species of Petrochromis

### ***Petrochromis calliris*** sp. nov

**Holotype**: NRM 57979, male 160.1 mm SL, Tanzania, Cape Mpimbwe, S 7°10969, E 30°50227, rocky bottom with sand patches at a depth of 5 m, net, 2 Feb 2008, M. Karlsson.

**Paratypes**: NRM 57978-57979, 2 males and 2 females, 156.8-176.8 mm SL; NRM 51319, male, 179.0 mm SL; same data as holotype.

**Diagnosis**: Distinguished from other *Petrochromis* by 1) greater number of vertebrae (33-34)compared to all other *Petrochromis* (30-33) except *P. heffalumpus* (34-35), 2) greater number of gill rakers on lower limb of first arch (13-15) compared to all other *Petrochromis* (10-14) except *P. orthognathus* (11- 15), 3) dorsal and anal fins in males have blue coloring near the edges.

D. XVIII-XIX, 9-10; A. III, 7-8; P. 14; gill-rakers on lower limb of first gill arch 13-15; lateral-line scales 34-35; vertebrae 33-34 (usually 33).

Body moderately deep. Snout profile of ascending processes of premaxillae moderately prominent. Upper jaw extending in front of lower jaw notably. Posterior margins of lip not reaching to vertical from anterior orbital rim. Tricuspid teeth on jaws in high density, not showing regular series. Cheek with 5-6 rows of scales. Scales present on opercle. Small scales present on base of dorsal fin. Last dorsal spine and 4th soft ray longest. Third anal spine and 4th soft ray longest. Posterior tips of dorsal and anal soft rays slightly filamentous. Pectoral fin shorter than head. Pelvic fin reaching first anal spine. Caudal fin subtruncate.

Color in life. Body color brown to green, often with blue and yellow areas. Fins tend to be reddish and to have blue coloring near the edges. Caudal fin often dark brownish.

Color in preservative. Body and fins dark brown.

**Habitat and feeding behaviour**: *Petrochromis calliris* occurs between Udachi and Kala Bay, usually at depths of 1-10 m. It is very territorial and males defend territories in shallow rocky areas. Usually it does not inhabit the same areas as *P. macrognathus*. *Petrochromis calliris* does not live quite as shallow as *P. macrognathus* but nevertheless the two species seem to compete for resources. *Petrochromis calliris* scrape algae off stones and rocks for food.

**Remarks**: *Petrochromis calliris* is similar to *P. heffalumpus* in the number of dorsal-fin spines (XVIII- XIX); the more numerous vertebrae, and the subtruncate caudal fin.

In the ornamental fish trade *P. calliris* is referred to as *Petrochromis* sp. Kasumbe rainbow.

### ***Petrochromis daidali*** sp. nov

**Holotype**: NRM 58000, male 159.7 mm SL, Tanzania, Nkwasi Point, S 6°25447, E 29°73257, rocky bottom with sand patches at a depth of 5 m, net, 17 Dec 2007, M. Karlsson.

**Paratypes**: NRM 58000, 2 males and 1 female, 129.3-153.1 mm SL, same data as holotype. NRM 57983, 4 males, 147.0-189.7 mm SL; NRM 51318, male, 207.1 mm SL; Tanzania, Cape Mbimbwe, 7.10969°S, 30.50227°E, 2 Feb 2008, M. Karlsson. NRM 51287, 5 males and 3 females, 139.8-187.0 mm SL; NRM 60212, female, 197.3 mm SL; Tanzania, Kansombo, M. Karlsson.

**Diagnosis**: Distinguished from other *Petrochromis* by 1) a slightly increased number of lateral-line scales (34-36) compared to all other *Petrochromis* (30-35) except *P. heffalumpus* (35-38) and *P. trewavasae* (32-36), 2) a slightly increased number of pectoral rays (14-15) compared to all other *Petrochromis* (13-14) except *P. heffalumpus* (14-15) and *P. macrognathus* (15), 3) males have a labyrinth pattern on their head.

D. XVIII-XIX, 9-10; A. III, 7-8; P. 14-15; gill-rakers on lower limb of first gill arch 11-14; lateral-line scales 34-36; vertebrae 32-33 (usually 33).

Body deep. Snout profile of ascending processes of premaxillae moderately prominent. Upper jaw extending in front of lower jaw notably. Posterior margins of lip not reaching to vertical from anterior orbital rim. Tricuspid teeth on jaws in higher density, not showing regular series. Cheek with 4-5 rows of scales. Scales present on opercle. Last dorsal spine and 4th soft ray longest. Third anal spine and 3rd soft ray longest. Posterior tips of dorsal and anal soft rays not filamentous. Pectoral fin shorter than head. Pelvic fin reaching first anal spine. Caudal fin emarginate.

Color in life. *Petrochromis daidali* have many colormorphs but generally males have a blue body and also a labyrinth pattern on their head. Brown and green color varieties can also be found. Females have less conspicuous colors than males, often greyish green. In different locations you can find different shades of blue amongst the male populations. In Cape Mpimbwe they are steely blue, in Kipili they are dark blue, in Mtosi they are shining blue, in Ninde they are blue with whitish spots on the back and at Izinga Island they have a blue head and a yellow body.

Color in preservative. Body and fins greyish brown. Face pattern still visible although slightly faded. **Habitat and feeding behaviour**: *Petrochromis daidali* are usually found at depths of 2-10 m. Both sexes are fiercely territorial and prefer a rocky environment with stones and boulders. Especially males continually patrol their territory which is a 15-25m^2^ large area which often includes at least some caves. In shallower waters the territories are smaller and the number of competing males increases. *Petrochromis daidali* scrape algae off stones and boulders for food.

**Remarks**: *Petrochromis daidali* is similar to *P. polyodon* in morphometry (Table 4) and Karlsson & Karlsson (2008) inform me that *P. daidali* is sometimes mistaken for *P. polyodon*. The most conspicuous thing about *P. daidali* is the unique labyrinth-like face pattern present on most males. In the ornamental fish trade *P. daidali* is referred to as *Petrochromis* sp. Texas.

**Table 4.** Proportional measurements as the percent of standard length (SL) in species of *Petrochromis* Data show ranges, means are in parenthesis

|  | *P.polyodon* | *P.fasciolatus* | *P.trewavasae* | *P.orthognathus* | *P.famula* | *P.macrogynathus* |
| --- | --- | --- | --- | --- | --- | --- |
| # Specimens | 9* | 15 | 3** | 23*** | 9 | 6 |
| Standard length (mm) | 113.3-170.3 | 86.7-130.7 | 111.0-128.2 | 93.0-128.2 | 97.3-136.2 | 89.8-150.0 |
| Head length | 33.1-38.7 (35.3) | 30.8-34.7 (32.7) | 32.9-36.3 (34.5) | 31.9-35.6 (33.4) | 34.3-38.2 (36.2) | 34.6-35.9 (35.5) |
| Snout length | 18.0-20.1 (19.1) | 14.2-16.3 (14.8) | 17.4-19.5 (18.5) | 15.1-17.6 (16.1) | 17.9-20.0 (18.9) | 16.5-19.5 (18.5) |
| Snout width | 17.1-19.3 (18.4) | 14.9-16.7 (16.0) | 16.8-20.5 (18.5) | 13.4-16.2 (15.0) | 17.1-21.5 (18.7) | 15.6-19.2 (17.9) |
| Depth of body | 37.3-41.5 (39.1) | 34.6-38.9 (37.0) | 38.2-42.1 (39.9) | 34.2-40.9 (37.2) | 37.9-42.0 (40.3) | 33.9-36.4 (35.5) |
| Eye diameter | 7.3-8.8 (8.1) | 7.3-9.8 (8.4) | 8.3-9.4 (8.8) | 8.0-9.9 (8.7) | 7.1-8.5 (7.8) | 7.0-9.0 (7.7) |
| Interorbital width | 10.9-12.8 (11.6) | 10.7-12.2 (11.4) | 11.3-11.8 (11.6) | 9.5-12.1 (10.7) | 12.2-13.4 (12.8) | 10.1-11.2 (10.6) |
| Pectoral fin length | 28.8-34.0 (32.0) | 23.3-33.5 (28.5) | 30.1-32.3 (31.4) | 30.0-35.0 (32.5) | 34.6-38.1 (36.0) | 25.5-29.3 (27.6) |
| Length of upper jaw | 13.7-16.6 (15.4) | 10.4-13.6 (12.5) | 12.2-14.9 (13.7) | 11.1-13.9 (12.6) | 13.5-17.0 (15.7) | 13.4-16.0 (14.9) |
| Length of lower jaw | 12.0-14.0 (13.0) | 10.6-15.1 (12.5) | 10.8-14.0 (12.2) | 10.6-12.9 (11.9) | 12.1-15.6 (13.8) | 11.1-12.5 (11.9) |
| Labial depth | 3.8-4.8 (4.4) | 2.8-3.8 (3.3) | 4.1-4.4 (4.3) | 2.8-4.3 (3.6) | 4.6-5.1 (4.8) | 4.4-4.8 (4.6) |
| Depth of caudal peduncle | 11.0-12.6 (11.8) | 10.5-11.6 (11.0) | 10.8-12.2 (11.4) | 9.2-11.5 (10.5) | 11.6-12.4 (12.0) | 11.0-12.7 (11.6) |
| Length of caudal peduncle | 15.8-18.3 (17.0) | 14.1-17.3 (15.9) | 10.8-16.3 (14.1) | 14.4-17.5 (15.9) | 14.0-15.8 (15.1) | 16.4-17.6 (16.9) |
| Length of last dorsal spine | 13.2-16.8 (14.4) | 11.1-15.2 (13.3) | 14.6-15.0 (14.8) | 12.2-15.0 (13.6) | 13.6-15.6 (14.6) | 12.1-14.3 (13.2) |
|  | *P.ephippium* | *P.calliris* | *P.daidali* | *P.heffalumpus* | *P.lisachisme* | *P.paucispinis* |
| # Specimens | 8 | 6 | 18 | 7 | 12 | 4 |
| Standard length (mm) | 127.5-156.9 | 153.2-179.0 | 129.3-207.1 | 195.4-281.2 | 133.9-202.8 | 128.7-156.3 |
| Head length | 33.6-36.3 (34.6) | 32.6-36.2 (32.7) | 32.7-37.7 (35.1) | 33.5-35.7 (34.4) | 32.0-38.3 (34.3) | 35.2-37.6 (36.1) |
| Snout length | 17.1-18.6 (17.7) | 17.5-18.7 (18.0) | 17.4-19.6 (18.7) | 18.4-19.8 (18.9) | 16.0-18.4 (16.9) | 18.3-20.6 (19.3) |
| Snout width | 16.8-20.6 (18.6) | 16.3-17.8 (16.9) | 16.9-21.9 (19.2) | 17.3-19.1 (17.8) | 16.7-19.7 (18.1) | 19.6-22.0 (20.7) |
| Depth of body | 37.2-41.1 (39.4) | 36.5-39.9 (38.5) | 35.8-41.5 (38.8) | 38.9-45.4 (41.7) | 39.1-44.3 (41.1) | 40.8-46.1 (43.2) |
| Eye diameter | 7.8-9.4 (8.4) | 6.5-8.1 (7.2) | 6.5-8.5 (7.5) | 6.1-7.1 (6.6) | 6.8-9.6 (7.7) | 7.8-8.3 (8.0) |
| Interorbital width | 12.2-13.2 (12.6) | 11.7-12.9 (12.2) | 11.3-14.2 (12.3) | 12.4-15.0 (13.6) | 11.7-13.3 (12.4) | 12.2-14.4 (13.2) |
| Pectoral fin length | 29.0-31.6 (30.2) | 29.8-33.4 (31.5) | 27.8-33.0 (31.0) | 28.0-31.6 (30.0) | 28.5-32.9 (30.0) | 30.3-34.8 (32.7) |
| Length of upper jaw | 13.4-15.2 (14.2) | 13.2-15.1 (14.3) | 13.7-17.7 (15.3) | 13.3-17.3 (14.9) | 13.8-15.9 (14.8) | 14.6-17.1 (15.7) |
| Length of lower jaw | 11.3-13.4 (12.3) | 12.2-13.8 (12.9) | 12.0-15.2 (13.3) | 11.4-16.4 (13.6) | 11.4-13.7 (12.5) | 13.2-14.4 (13.6) |
| Depth of labial | 3.0-4.2 (3.7) | 3.5-4.4 (4.0) | 3.7-4.5 (4.1) | 3.4-4.3 (4.0) | 2.9-4.2 (3.5) | 3.8-4.6 (4.2) |
| Depth of caudal peduncle | 10.5-12.5 (11.6) | 11.2-12.5 (11.8) | 11.3-13.4 (12.0) | 11.4-16.4 (13.2) | 11.5-12.5 (12.1) | 11.9-13.4 (12.6) |
| Length of caudal peduncle | 14.4-17.3 (15.8) | 16.2-17.4 (16.7) | 15.7-18.4 (17.1) | 16.7-18.7 (17.4) | 14.8-18.2 (16.5) | 16.2-17.4 (16.7) |
| Length of last dorsal spine | 13.5-15.6 (14.4) | 12.2-14.4 (13.5) | 11.8-16.6 (13.6) | 11.9-14.7 (13.2) | 12.2-16.0 (13.9) | 13.5-14.4 (13.9) |

\*5 juveniles excluded; \*\*1 juvenile and 13 aquarium specimens excluded; \*\*\*4 aquarium specimens excluded

### ***Petrochromis heffalumpus*** sp. nov

**Holotype**: NRM 57992, male 281.6 mm SL, Tanzania, Cape Mpimbwe, S 7°10969, E 30°50227, rocky bottom with sand patches at a depth of 15 m, net, 2 Feb 2008, M. Karlsson.

**Paratypes**: NRM 57989-57991, 3 females, 195.4-235.8 mm SL; NRM 57993-57994, 2 males, 263.9-269.2 mm SL; NRM 51316, male, 281.2 mm SL; same data as holotype.

**Diagnosis**: Distinguished from other *Petrochromis* by 1) greater size (195.4-281.6 mm SL instead of<209.8 mm SL), 2) greater number of lateral-line scales (35-38 instead of 30-36), 3) greater number of vertebrae (34-35 instead of 30-34), 4) greater number of dorsal soft rays (10-11 instead of 8-10), 5) caudal fin flattened horizontally on top and bottom.

D. XVIII-XIX, 10-11; A. III, 8-9; P. 14-15; gill-rakers on lower limb of first gill arch 11-12; lateral-line scales 35-38; vertebrae 34-35 (usually 34).

Body deep, especially in male specimens. Snout profile slightly humped by ascending processes of premaxillae. Upper jaw extending in front of lower jaw notably. Posterior margins of lip not reaching to vertical from anterior orbital rim. Tricuspid teeth on jaws in high density, not showing regular series. Cheek with 5-6 rows of scales. Scales present on opercle. Small scales present on bases of dorsal and anal fins. Last dorsal spine and 5th soft ray longest. Third anal spine and 4th soft ray longest. Posterior tips of dorsal and anal soft rays not filamentous. Pectoral fin shorter than head. Pelvic fin reaching anal spines, even reaching slightly beyond the anal spines in larger male specimens. Caudal fin subtruncate.

Color in life. Body brown to blue. Fins tend to be blue and become darker further out from the body.

Color in preservative. Body and fins dark brown.

**Habitat and feeding behaviour**: *Petrochromis heffalumpus* occurs between Lyamembe and Izinga island, usually at depths of 10-25 m. It prefers an environment with large boulders and sand patches. Males are fiercely territorial and reside near a large cave that they can protect. *Petrochromis heffalumpus* scrape algae off large flat stony surfaces.

**Remarks**: *Petrochromis heffalumpus* share a number of characteristics with *P. macrognathus*. It has a similar number of dorsal spines, dorsal soft rays, pectoral rays and anal soft rays. The snout profile of *P. heffalumpus* is not quite as pronounced as *P. macrognathus* but nevertheless shows a similar hump. *Petrochromis heffalumpus* differs from *P. macrognathus* in that it has a deeper body. *Petrochromis macrognathus* also has a proportionally wider snout, fewer vertebrae and fewer lateral-line scales. *Petrochromis heffalumpus* is also similar to *P. calliris* in that they have a similar number of dorsal spines and vertebrae and also they both have a subtruncate caudal fin. The body proportions of these two species are also closer to each other than if you compare them to *P. macrognathus*.

In the ornamental fish trade *P. heffalumpus* is referred to as *Petrochromis* sp. Blue giant.

### ***Petrochromis lisachisme*** sp. nov

**Holotype**: NRM 57973, male 194.3 mm SL, Tanzania, Lyamembe, 6.46771°S, 29.93698°E, rocky bottom with sand patches at a depth of 35 m, net, 19 Jan 2008, M. Karlsson.

**Paratypes**: NRM 57995-57996, 4 males and 2 females, 133.9.5-187.4 mm SL; NRM51315, male, 176.8 mm SL; Tanzania, Cape Mbimbwe, 7.10969°S, 30.50227°E, 2 Feb 2008, M. Karlsson. NRM 57973-57974, 4 males, 194.1-209.8 mm SL, same data as holotype.

**Diagnosis**: Distinguished from other *Petrochromis* by 1) greater number of dorsal spines (XVIII-XXI instead of XVII-XX), 2) having a lunate caudal fin (*P. trewavasae* shares this trait), 3) males have a distinct forehead which also gives the jaws a protruded appearance.

D. XVIII-XXI, 8-10; A. III, 7-8; P. 13-14; gill-rakers on lower limb of first gill arch 12-13; lateral-line scales 34-35; vertebrae 32-33.

Body deep. Snout profile slightly humped by ascending processes of premaxilla. Upper jaw prognathous. Posterior margins of lip not reaching to vertical from anterior orbital rim. Tricuspid teeth on jaws in low density, showing regular series. Cheek with 4-6 rows of scales. Scales present on opercle. Small scales present on base of dorsal fin. Last dorsal spine and 4th soft ray longest. Third anal spine and 4th soft ray longest. Posterior tips of dorsal and anal soft rays slightly filamentous. Pectoral fin shorter than head. Pelvic fin reaching first anal spine or slightly beyond. Caudal fin lunate.

Color in life. Body ranging from intense red, with a black patch around the eye and chin; to orange, occasionally with yellow spots on the fins; to greyish green. Fins in the intense red variety tend to be dark whereas in other color varieties the fins tend to be the same color as the body.

Color in preservative. Specimens from Cape Mpimbwe; body and fins greyish brown. Specimens from Lyamembe; body brown and fins dark brown, head with dark brown face and a light grey patch around the eye and chin.

**Habitat and feeding behaviour**: *Petrochromis lisachisme* occurs between Bulu Point and Kala Bay, usually at depths of 20-35 m. Adults are found only below 20 m and larger males, 20-25 cm, only below 35 m. It prefers an environment with boulders and sand patches. It is less territorial than most other *Petrochromis* and they patrol a relatively large area which often includes at least some caves. The spawning area is created against a large rock in the form of a half crater in the sand. *Petrochromis lisachisme* scrape algae off stony surfaces and also eat phytoplankton.

**Remarks**: *Petrochromis lisachisme* is similar to *Petrochromis trewavasae* in having long filamentous fins and a lunate caudal fin. They are also the only two *Petrochromis* with as many as 20 dorsal spines. Field observations suggest that the red color of *P. lisachisme* is more intense in more northern localities. The most intense red color is present in specimens from Lyamembe and Bulu point. Specimens from Cape Mpimbwe are orange, and in Kala Bay specimens are greyish.In the ornamental fish trade *P. lisachisme* is referred to as *Petrochromis* sp. Red.

### ***Petrochromis paucispinis*** sp. nov

**Holotype**: NRM 57971, male 156.3 mm SL, Tanzania, Halembe, S 5°73712, E 29°92167, rocky bottom with sand patches at a depth of 5 m, net, 8 Jan 2008, M. Karlsson.

**Paratypes**: NRM 57971, 2 females and 1 male, 128.7-144.8 mm SL, same data as holotype.

**Diagnosis**: Distinguished from other *Petrochromis* by 1) fewer number of dorsal spines (XVII) compared to all other *Petrochromis* (XVIII-XXI) except *P. polyodon* (XVII-XIX) and *P. famula* (XVII-XVIII), 2) fewer number of dorsal spines and rays combined (26) compared to most other *Petrochromis* (generally 27-29) except *P. polyodon* (26-28), 3) greater snout width (mean 20.7 % of SL) compared to all other*Petrochromis* species (with means of 15.0-19.2 % of SL).

D. 17, 9; A. III, 7; P. 14; gill-rakers on lower limb of first gill arch 11-12; lateral-line scales 34; vertebrae 31-32 (usually 31).

Body deep. Snout profile of ascending processes of premaxillae moderately prominent. Upper jaw extending in front of lower jaw notably. Posterior margins of lip almost reaching to vertical from anterior orbital rim. Tricuspid teeth on jaws in high density, not showing regular series. Cheek with 4- 6 rows of scales. Scales present on opercle. Last dorsal spine and 4th soft ray longest. Third anal spine and 3rd soft ray longest. Posterior tips of dorsal and anal soft rays not filamentous. Pectoral fin shorter than head. Pelvic fin reaching first anal spine. Caudal fin emarginate.

Color in life. Body blue and green with orange patches, the ventral part silvery, in males. Body green with vague orange patches in females. Fin color orange in both sexes.

Color in preservative. Body and fins greyish brown.

**Habitat and feeding behaviour**: *Petrochromis paucispinis* occurs between Karilani Island to Mayobozi, usually at depths of 1-15 m. Both sexes are fiercely territorial and are especially aggressive towards other *P. paucispinis*. Territories are 10-20m^2^, consist of rocky caves and are usually in shallower water than 10m. *Petrochromis paucispinis* scrape algae off stones for food.

**Remarks**: *Petrochromis paucispinis* is similar to *P. polyodon* and all seven meristic characters overlap (Table 5). *Petrochromis paucispinis* tend to have slightly fewer dorsal spines, vertebrae and gill rakers on the lower limb of the first arch, but often has more lateral-line scales than *P. polyodon*. In the ornamental fish trade *P. paucispinis* is referred to as *Petrochromis* sp. Kasumbe.

**Table 5.**
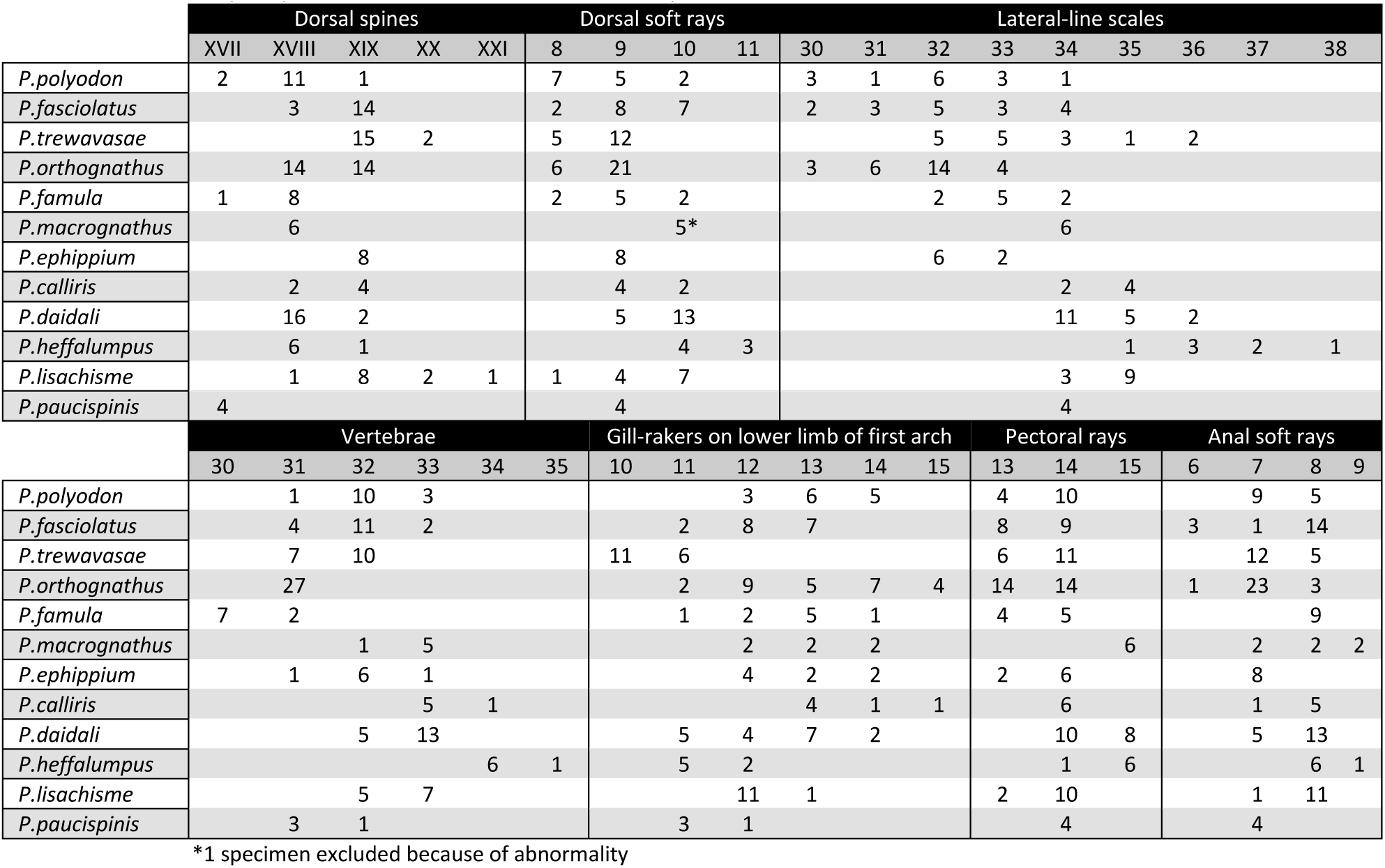
Frequency distribution of 7 meristic characters in species of *Petrochromis*

**Table 6.** Key to the species of Petrochromi

| Table 6. Key to the species of *Petrochromis* |  |  |
| --- | --- | --- |
| 1a | Jaws equal anteriorly, or upper jaw projecting slightly | 2 |
| 1b | Upper jaw projecting in front of lower jaw notably | 3 |
| 1c | Lower jaw projecting | *P. fasciolatus* |
| 2a | Upper jaw projecting slightly; Body deep; Tricuspid teeth in dense pad; Cheek scales not present on lower part of cheek | *P. famula* |
| 2b | Jaws equal anteriorly; Body not so deep; Tricuspid teeth not in a dense pad; Teeth showing regular series | *P. orthognathus* |
| 3a | Caudal fin subtruncate | 4 |
| 3b | Caudal fin lunate | 5 |
| 3c | Caudal fin emarginate | 6 |
| 4a | Body color brown to green; Fin color often reddish; Body not so deep; Gill-rakers on lower limb of first arch 13-15 | *P. calliris* |
| 4b | Body color brown to blue; Fin color blue; Body deep; Gill-rakers on lower limb of first arch 11-12 | *P. heffalumpus* |
| 5a | Body color dark brown; Gill-rakers on lower limb of first arch 10-11; Tricuspid teeth in a dense pad | *P. trewavasae* |
| 5b | Body color intense red to greyish orange; Gill-rakers on lower limb of first arch 12-13; Tricuspid teeth not in a dense pad | *P. lisachisme* |
| 6a | Body deep; Convexity of premaxillary ascending processes on snout less pronounced; No small scales present on bases of dorsal and anal fins | 7 |
| 6b | Body not so deep; Convexity of premaxillary ascending processes on snout much pronounced; Many small scales present on bases of dorsal and anal fins; Pronounced concavity of chin region | *P. macrognathus* |
| 7a | Caudal fin emarginate, but closer to lunate than subtruncate; Snout wide, often >19 % of SL; At least 9 dorsal soft rays | 8 |
| 7b | Caudal fin emarginate, but closer to subtruncate than lunate; Snout not so wide, often <19 % of SL; As few as 8 dorsal soft rays | *P. polyodon* |
| 8a | Dorsal spines 18-19; Dorsal spines and rays combined 27-28 | 9 |
| 8b | Dorsal spines 17; Dorsal spines and rays combined 26; Body colored with orange and green, males also with blue | *P. paucispinis* |
| 9a | Lateral-line scales 34-36; Labyrinth pattern on face (♂) | *P. daidali* |
| 9b | Lateral-line scales 32-33; Body color brown to yellow-orange; Saddle-like rectangular light-coloured patch under dorsal fin | *P. ephippium* |

### Phylogenetic analysis

The morphological matrix (Table 3) yielded a strict consensus tree (Fig. 5) with *P. orthognathus, P. fasciolatus* and *P. famula* as the three species most closely related to the outgroup *Tropheus*. No significant support could be found for this tree. One character needing further explanation is the correlation between vertebrae and lateral-line scales (character 8 in Table 3). Working with the hypothesis that all *Petrochromis* would have approximately the same number of vertebrae (V) as lateral-line scales (L) I correlated these two for each specimen (Table 7). V*min* is the least amount of lateral-line scales compared to the number of vertebrae, on any single specimen, within a species, V*max* being the opposite. The results show that most species have more lateral-line scales than vertebrae, excluding some specimens of *P. polyodon, P. fasciolatus* and *P. orthognathus*, and that no species has a V*min* value below -2 or a V*max* value of more than 4. This can be described as [V=(L+1)±3].

**Fig. 5.**
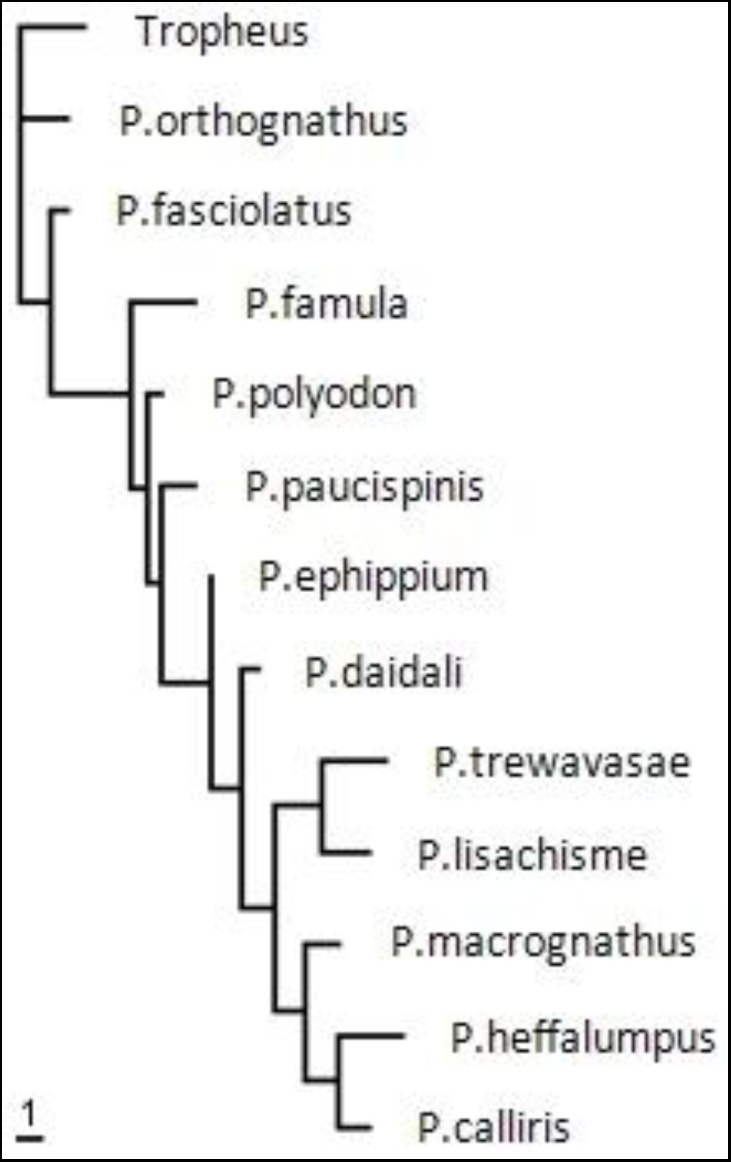
Strict consensus tree of *Petrochromis* based on morphological data and parsimony analysis.

**Table 7.** Number of vertebrae (V) compared to the number of lateral-line scales

|  | *Vmin* | *Vmax* |
| --- | --- | --- |
| *P. polyodon* | -2 | 3 |
| *P. fasciolatus* | -2 | 2 |
| *P. trewavasae* | 0 | 4 |
| *P. orthognathus* | -1 | 2 |
| *P. famula* | 1 | 4 |
| *P. macrognathus* | 1 | 2 |
| *P. ephippium* | 0 | 1 |
| *P. calliris* | 1 | 2 |
| *P. daidali* | 1 | 4 |
| *P. heffalumpus* | 1 | 4 |
| *P. lisachisme* | 1 | 3 |
| *P. paucispinis* | 2 | 3 |

The Maximum Likelihood and Bayesian trees created in the molecular analyses are shown in Figure 6. The cytochrome *b* analyses favour *P.* sp. Yellow as the sister group of all other *Petrochromis*, and *P. famula* as the sister group of all *Petrochromis* excluding *P.* sp. Yellow. In the d-loop analyses both trees favour *P. famula* and *P. orthognathus* as the sister group of all other *Petrochromis*. Also the d-loop Maximum Likelihood tree favours *P. fasciolatus* as the sister group of all other *Petrochromis*excluding *P. famula* and *P. orthognathus*.

**Fig. 6.**
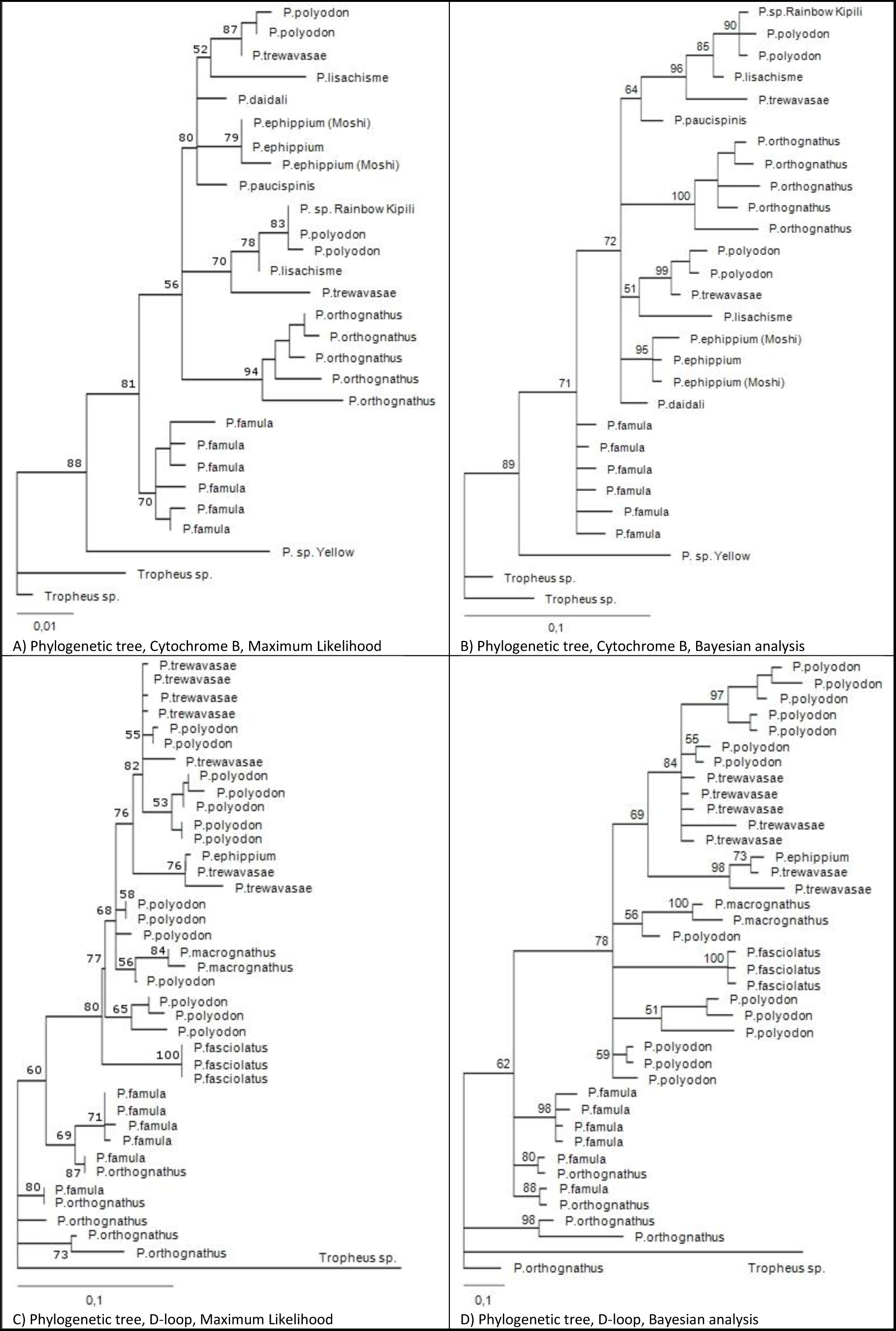
Hypotheses of phylogenetic relationships among species of *Petrochromis*, based on mitochondrial cytochrome *b* and d-loop sequences, subject to Maximum Likelihood and Bayesian inference analyses.

Combined the morphological tree and the four molecular trees place *P. orthognathus* and *P. famula* (clade *A*) as the sister group of all other *Petrochromis*. Some support is also found that *P. fasciolatus* (clade *B*) is the sister group of all other *Petrochromis* excluding *P. famula* and *P. orthognathus* (clade *C*). *P*. sp Yellow is here excluded due to insufficient data.

## Discussion

Based on phylogenetic analyses *Petrochromis* can be divided into three clades (Fig. 7). Clade *A* consisting of *P. famula and P. orthognathus*, clade *B* consisting of *P. fasciolatus* and clade *C* consisting of all other *Petrochromis*. This is similar to Yamaoka’s attempt at grouping together different species of *Petrochromis* based on the number of times a fish opens and closes its mouth per second and per feeding behaviour (Fig. 2) and also fits well with Sturmbauer’s molecular tree (Fig. 2). Further study of *Petrochromis* sp. Yellow is needed to determine its phylogenetic position but molecular results suggests it belongs in clade *A* (Fig. 6, A-B). Images by Herrmann (1994) show that it has isognathous jaws consistent with clade *A*. The three clades are also recovered in number of vertebrae and jaw position (Fig. 7). A trend towards an increased number of vertebrae in cichlids was suggested by Cichocki (1976) and seems to be the trend also within the genus *Petrochromis*.

**Fig. 7.**
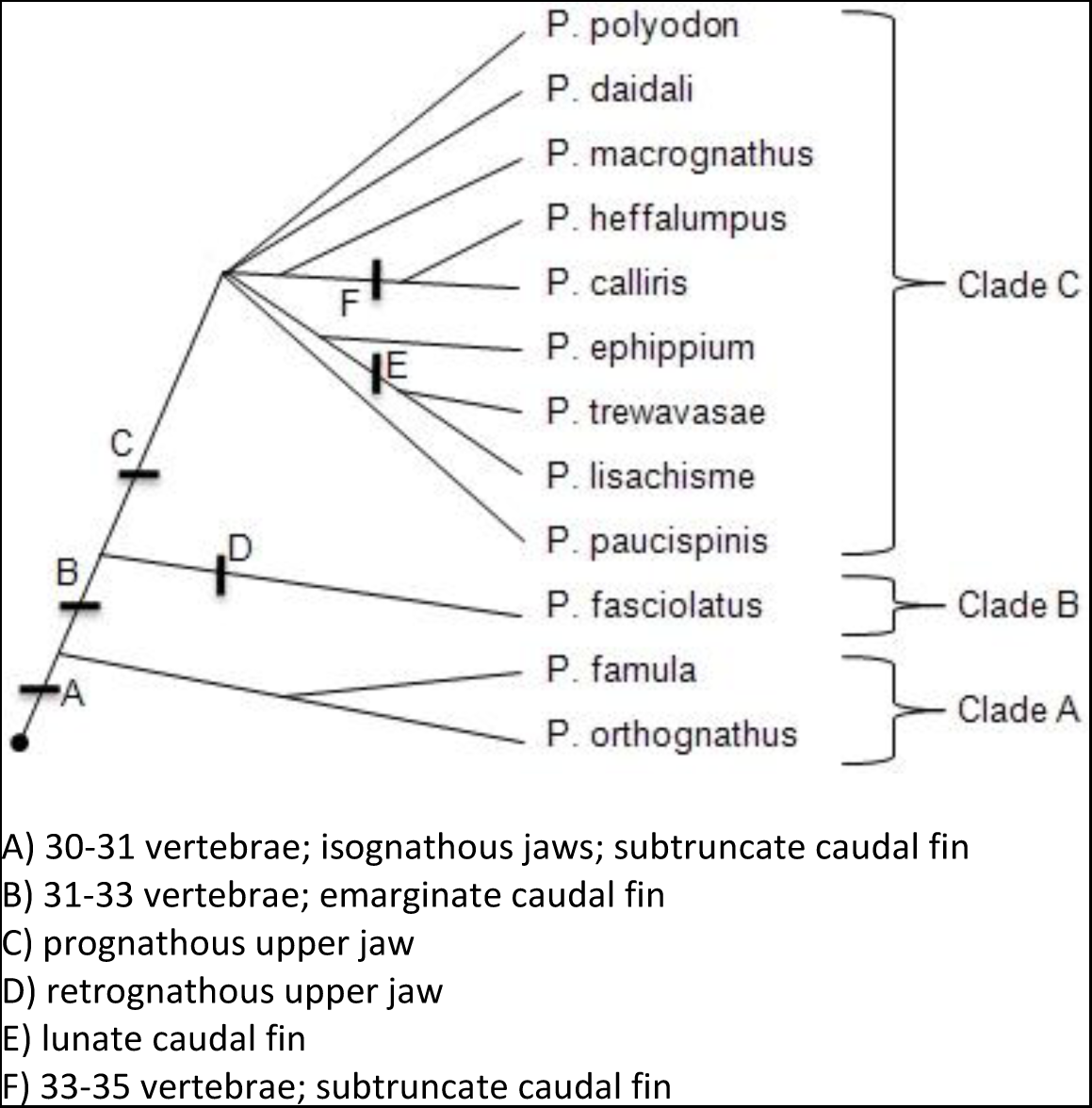
Hypothesis of the phylogeny of the genus *Petrochromis* based on morphological and molecular data.

The morphological tree (Fig. 5) suggests placing *P. fasciolatus* between *P. orthognathus* and *P. famula*, contrasting with molecular results, but also similar to Yamaoka’s morphological attempt at grouping together different species of *Petrochromis* based on number of jaw teeth and development of the adductor mandibulae muscles (Fig. 2). The morphological matrix analysis fails to take into account the unique retrognathous upper jaw in *P. fasciolatus*. No significant support could be found for the morphological tree but it is very similar to the molecular trees. The lack of support is probably mainly due to an insufficient amount of data.

The low resolution in clade *C* in the molecular analysis (Fig. 6) nevertheless holds points of interest. The sequences of *P. ephippium* and *P*. sp. Moshi are identical and they are of the same species (Fig. 6, A-B). In the trees based on d-loop (Fig. 6, C-D) the single *P. ephippium* groups with two specimens of *P. trewavasae* from GenBank, suggesting that the identification in GenBank is incorrect. It is noteworthy that two of the new species, *P. daidali* and *P. paucispinis*, fail to group with any other *Petrochromis* species within clade *C*.

Morphological data (Fig. 5) suggests a clade within clade *C* comprised of *P. calliris, P. heffalumpus* and *P*. *macrognathus*. That a deep living species like *P. heffalumpus* shares so many character states with *P. macrognathus*, which lives in shallower water is interesting. Closer inspection reveals remarkable similarities when it comes to fin ray counts (Table 5). The third species in this clade, *P. calliris*, is the most ordinary looking *Petrochromis* amongst the three. It lives shallower than *P. heffalumpus* but deeper than *P. macrognathus* (Table 1). Unfortunately no tissue was available of *P. calliris* or *P. heffalumpus* and these two species could therefore not be included in the molecular analyses. Another clade within clade *C*, also based on morphological results, is comprised of *P. trewavasae* and *P. lisachisme*, the only two species with lunate caudal fins. Yet again the pattern is that a deep-water species, *P. lisachisme*, shares many characters with a particular shallow-water species, *P. trewavasae*.

Basal tree positions (Figs. 5-7) suggest that the “ancestral *Petrochromis* “might have looked something like *P. orthognathus* (Fig. 8) with isognathous jaws and a relatively elongate body. That the teeth of *P. orthognathus* show regular series indicate that it is less adapted for epilithic algal feeding than e.g. *P. famula* (Yamaoka, 1982), and may also be an ancestral character. The condition is similar to the outgroup *Tropheus* which also has teeth showing regular series. The only species of *Petrochromis* except *P. orthognathus* showing regular tooth series is *P. lisachisme*, which is positioned higher up the morphological tree (Fig. 7), but in this species it is probably a reversal reflecting its deeper habitat and its more varied diet not limited to algae. Perhaps *Petrochromis* sp. Yellow shares this trait also, being placed so close to *Tropheus* in the phylogeny based on cytochrome *b* (Fig. 6, A-B), and having isognathous jaws. This however is speculatory since I only worked with its DNA and photographs of this potentially new species.

**Fig. 8.**
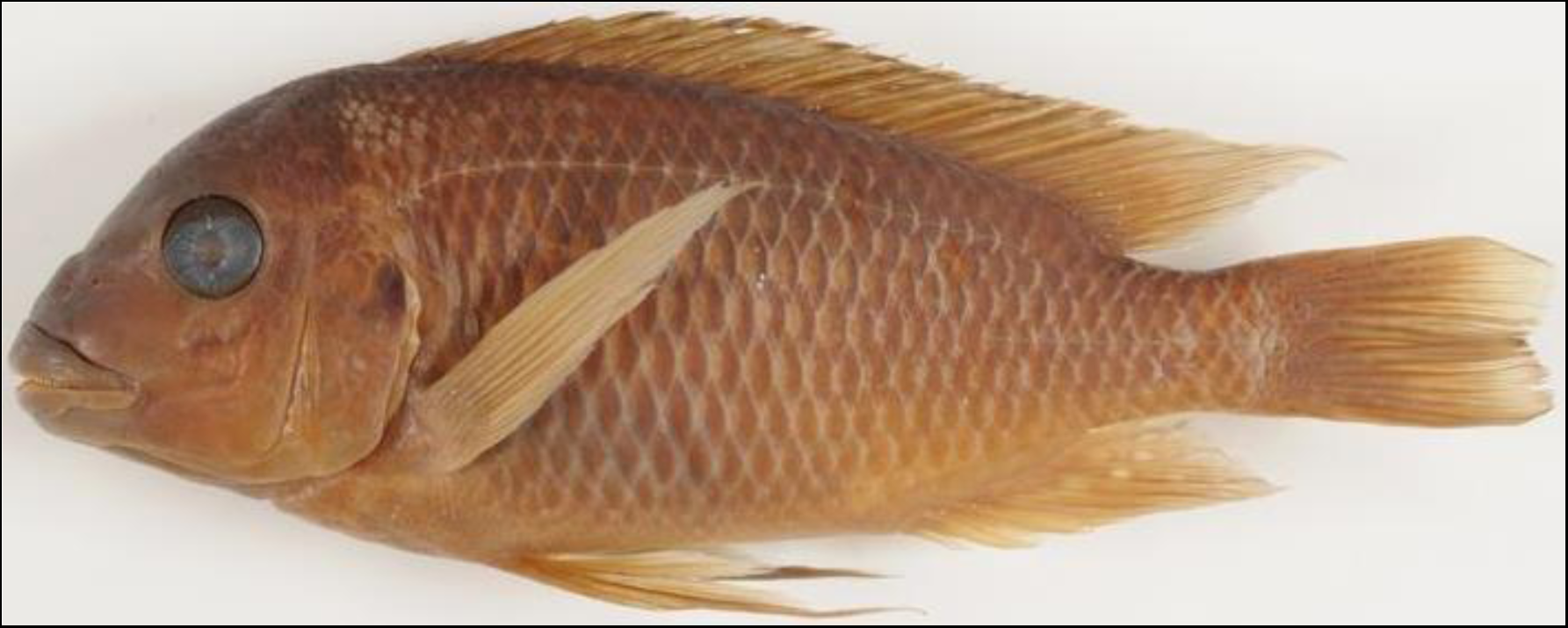
*Petrochromis orthognathus* (NRM 17447)

## Acknowledgements

I wish to thank Sven O. Kullander, Bo Delling and Erik Åhlander of the Swedish Museum of Natural History for their help during this study; I am further grateful to Elizaveta Mattsson, Anders Silfvergrip, Nicklas Wijkmark, Bodil Kajrup, Te Yu Liao, Omar Mechedal, Claes Dannbeck, Bodil Cronholm and Ismail Malikov for their assistance during this study; Magnus and Mikael Karlsson of the African Diving LTD, for supplying *Petrochromis* specimens and expedition data; Peter Nemeth for supplying tissue samples; Ulf Jondelius and Bertil Borg, professors of the Swedish Museum of Natural History and Stockholm University respectively, for reviewing this article; and a special thanks to the late Fang Kullander and Birgitta Tullberg.

